# Evaluation of salinity tolerance of lowland rice genotypes at the reproductive stage

**DOI:** 10.1101/2022.08.22.504861

**Authors:** Safidimanjato Rafaliarivony, Hery Lisy Tiana Ranarijaona, Mbolarinosy Rasoafalimanana, Tendro Radanielina, Matthias Wissuwa

## Abstract

Salinity is an abiotic stress considered as the most widespread soil problem next to drought in rice growing areas of the world. To facilitate agricultural production on salt-affected soil, water and soil management is the most common practice, but this approach is increasingly problematic because water is becoming scarce. Therefore, developing tolerant varieties would the best solution to this problem. The present study evaluated salinity tolerance of 72 lowland rice genotypes at the reproductive stage. Experiments were conducted in irrigated fields at Marovoay, Madagascar, under low and moderately high salinity with electric conductivities of 2 dSm^-1^ and 4 dSm^-1^, respectively. A subsequent validation experiment was conducted in a pot experiment at three levels of salinity corresponding to 0, 4 and 8 dSm^-1^. Plant height, panicle number, panicle length, panicle fertility, spikelet fertility, straw weight and grain yield were measured together with a visual score of salt injury. Field salinity strongly reduced panicle number and spikelet fertility, reducing grain yield to less than 10 g m^-2^ in sensitive genotypes compared to more than 60 g m^-2^ in tolerant genotypes. The field experiment classified 20% of genotypes as tolerant, 50% as intermediate and 30% as sensitive to salinity. Four genotypes IR55179, MTM13_1, MTM13_3, MTM13_5, were re-confirmed as highly tolerant in the pot experiment. Higher spikelet and panicle fertility in tolerant genotypes contributed to their superior grain yield under salinity stress whereas these traits were particularly reduced in the sensitive local varieties. Genotypes with tolerance to salinity at the reproductive stage identified here could be used as donors to improve grain yield of local sensitive varieties, possibly using spikelet and panicle fertility as selection criteria for screening breeding lines at the reproductive stage.

## Introduction

Salinity is an abiotic stress considered as the most widespread soil problem next to drought in rice growing areas of the world [1]. More than 800 million hectares of land throughout the world are adversely affected by high salinity [2]. Salinity affects about 50% of irrigated land worldwide which includes about 30% of the rice areas [3]. Salinity occurs when the accumulation of salts dissolved in solution of soil reach to a level that inhibits plant growth and development. It is characterized by the presence of excessive concentrations of soluble salts in the soil. The major species of salts are sodium chloride, magnesium and calcium sulphates and bicarbonates but the predominant is sodium chloride [4]. To quantify the degree of salinity, electric conductivity (EC) of either the saturation extracts or soil solution collected from the root zone are measured. A soil is characterized as saline if its EC is 4 dSm^-1^ or more [5]. To reduce the effect of salinity stress and maintain agricultural production on salt-affected soil, water and soil management are most common practices. Water management consists of leaching salts with fresh water through frequent irrigation, however, increasing water scarcity in many of the rice growing regions places some limitations on relying on this approach [6].

In Madagascar, the second most important rice granary is located in the north western coastal region of the country around the town of Marovoay [7]. High yielding varieties with a good response to fertilizer and adapted to the agro-ecosystem of the North Western zone have partly replaced traditional varieties characterized by late maturity and low yield potential [8]. Despite the potentiality of these varieties, agronomic yield remains low and shows great fluctuation of 1.7 to 2.9 tons per hectare [9]. Climate change has a dramatic effect on agricultural production in the region [10]. One factor behind reduced yield is salinity which occurs mainly throughout the dry season system and is caused by the rising sea water in the irrigated perimeters and hampered by water scarcity and increasing temperature [10,11].

Rice is generally characterized as a salt sensitive crop. Rice yield is affected by salinity at or above 3.0 dSm^-1^ and decreases 12% for every unit (dsm-1) increase in EC above 3.0 dSm^-1^ [12,13]. Rice yield in salt affected land is significantly reduced with an estimation of 30–50% yield losses annually [14]. The effect of salinity on rice depends on the severity of salinity, which in turn is affected by soil physical properties, water regime, crop growth stage, duration of exposure and temperature, humidity and solar radiation during exposure, as well as cultivar adaptations to these factors [15]. Rice is tolerant to salinity stress during germination and active tillering, whereas it displays higher sensitivity during early vegetative and reproductive stages [16]. It has been reported that salt tolerance at the reproductive stage is not correlated with tolerance at the seedling stage as they are controlled by different sets of genes [17,18]. Salt stress results in root growth inhibition, leaf rolling, and reductions in plant height, tiller number, panicle length and spikelet fertility. A delay in panicle emergence and flowering is also possible, which will lead to reduced yield in water-scarce environments [15,19]. Additionally, salt stress induces physiological and biochemical changes [20]. Several studies reported that the absorption and uptake of micro- and macro-mineral nutrients are altered under salinity stress, increased Na transport to the shoot and decreased K, Zn, and P uptake [20,21]. Likewise, it has been reported that salt stress increases ROS levels, causing significant injury and eventual death in plants [22].

Salinity stress induces metabolite changes; therefore, rice plants have evolved several physiological mechanisms to cope with salinity stress [2]. These mechanisms were classified into osmotic tolerance, ion exclusion, and tissue tolerance [23]. Osmotic tolerance concerns the adjustment of the osmotic potential in plant cells by accumulation of organic and/or compatible osmolytes such as proline, trehalose, and glycine betaine [24,25]. It has been reported that a salt tolerant rice cultivar accumulated more proline during salinity stress compared to salt sensitive rice cultivar [25]. Proline is an essential osmoregulator in plants exposed to hyperosmotic stresses such as drought and salinity [25]. Ion exclusion mainly involves Na+ and Cl–transport processes in roots, which prevent the excess accumulation of Na+ and Cl– in leaves [3]. The main seedling stage salinity tolerance QTL *Saltol* appears to be involved in maintaining shoot Na+/K+ homeostasis [25,26]. Production of enzymes catalyzing detoxification of reactive oxygen species, sequestration of Na+ in the vacuole and program cell death are involved in tissue tolerance [25]. Plant vigor, an avoidance mechanism, is also a major determinants of salt tolerance in rice [27]. Yeo et al (1990) [28] suggested that the concentration of transported Na+ will be lower in fast growing rice genotypes than in slowly growing ones.

Numerous studies have been undertaken to understand the different physiological and genetic factors associated with salinity tolerance of rice at the seedling stage and a main salinity tolerance QTL (*Saltol*) had been identified on chromosome 1 from donor variety Pokkali. However, only a few studies dealt on reproductive stage salinity tolerance [17,29]. Ahmadizadeh et al. (2016) [30] stated that there is a lack of reliable reproductive stage-specific phenotyping techniques and incomplete knowledge of the stage-specific mechanisms of salinity tolerance. Ahmadizadeh et al. (2016) [30] also specified that the complexity of the trait makes phenotyping for this trait exceedingly difficult, so that progress on reproductive stage salinity tolerance remains slow and still elusive. The present study is therefore contributing in solving problems linked with soil salinity by searching salinity tolerant genotypes at the reproductive stage. Specific objects of this study are (1) to evaluate the salinity tolerance of 72 lowland rice genotypes at the reproductive stage under field condition and validate the result in pot experiments, (2) to cluster the genotypes according to their reproductive stage salinity tolerance and (3) identify the most important agro morpho-physiological descriptors of salinity tolerance at the reproductive stage.

## Materials and methods

### Plant materials

Tolerance to salinity at the reproductive stage was evaluated in 72 lowland rice lines, of which 68 lines were selected from the *Pup1* breeding program in Madagascar. These lines were selected from a cross of two BC2 IR64-*Pup1* introgression lines, which in turn were derived from a cross of IR64 with *Pup1* donor line NIL14-4, which is a *Pup1* locus introgression line in the Nipponbare background (Wissuwa, unpublished). Progeny of the cross of two BC2 IR64-*Pup1* lines was imported from the Japan International Research Center for Agricultural Sciences (JIRCAS) to Madagascar in the F3 generation and selected for performance under low-input conditions in the F3 and F5 generations, with F4 and F6 generations being used for seed increases. The lines used in this experiment were in the F7 generation and are referred to as MTM lines. The remaining four lines used are local varieties obtained from National Center for Applied Research on Rural Development (FOFIFA).

### Field experiment

Field evaluation for salinity tolerance of the 72 lowland rice genotypes was carried out during the dry season of 2019 (May to September) under low and high salinity in two irrigated farmers’ fields located in the community of Marovoay, coastal North-Western Madagascar. The experiment was carried out following an augmented randomized complete block design, in which frequently repeated check varieties are presumed as fixed effects and are used to estimate position effects [31]. In this experiment, the design contained 10 blocks of ten genotypes, each block having 8 test entries and the two check entries IR64 and Tsipala A. Test entries were the 68 MTM lines plus4 local varieties. Four-week-old seedlings were transplanted as single seedlings/hill with a 20 cm x 20 cm spacing within and between rows. One plot consisted of a double row of 2 m length. No fertilizer was applied as is farmer’s practice in the region. Hand weeding were performed to control weeds. Pesticide was applied to control harmful bugs and rat attacks.

Low and high salinity fields were identified by interviewing local rice farmers, followed by electro-conductivity (EC) analyses using a handheld EC-meter. Homogeneous fields with reliable access to irrigation were chosen. At the point of testing the low salinity field had an EC of 2 dSm^-1^ whereas the severely saline field had an EC around 4 dSm^-1^. Salt stress was avoided between the early vegetative to maximum tillering stages by flushing salinity with frequent irrigation. When the majority of lines reached the maximum tillering stage, natural occurring salinity was induced by reducing irrigation intervals while maintaining a 5-10 cm standing water level. Salinity was monitored every two weeks during the growing period using an EC meter and irrigations were adjusted as needed to maintain target EC levels below 3 and above 4 dSm^-1^ for the low and severe salinity treatments, respectively.

### Validation experiment (pots)

To confirm observations and genotype rankings observed in the field, a pot culture experiment was carried out at the National Center for Applied Research on Rural Development (FOFIFA) in Mahajanga, Madagascar. The experiment was conducted during the end of rainy season to dry season (March to June) of 2020 and installed in an open area but covered with nets to protect from animals. Of the initial 72 lowland rice genotypes used in the 2019 field experiment, the eight most tolerant lines based on grain yield and yield related traits were tested in the pot experiment together with local check Sebota 281 (sensitive) and Tsipala A (tolerant). Salinity was induced during the reproductive stage only, following a protocol described by Gregorio et al. (1997) [32]. An experimental unit was a PVC tube segment of 15 cm height and 11 cm diameter that contained a cotton cloth bag filled with paddy field soil from Marovoay. Three of these PVC pots were placed in a big plastic pot (30 cm high and 30 cm diameter) filled with tap water which served as a water bath, simulating flooded field conditions. To facilitate the movement of water and solutes to the root zone, tubes were perforated with holes of 3 to 4 mm diameter drilled at a 2 cm distance. Tubes were furthermore open at the bottom. Fertilizer was applied as compound NPK (11-22-16) at a rate of 4 g/kg of soil at the time of filling cloth bags. Pots were arranged in a randomized complete block design (RCBD) with four replications, where a set of four big plastic pots was taken as a block. Five seedlings (one-week old) per entry were transplanted into each pot, but thinned into two plants per pot after two weeks. The water level was maintained at one cm above the soil. When rice plants reached the maximum tillering stage, all water in the water bath was siphoned out and exchanged with salinized water. In addition to a control (0 dsm^-1^), two levels of salinized water were prepared by dissolving 4.5 g and 9 g of table salt per liter of tap water to achieve medium (4 dSm^-1^) and high (8 dSm^-1^) salinity levels, respectively. Every two weeks, all water in a water bath was replaced by new salinized water to maintain the same salinity level. The level of flooding was maintained daily at 1 cm above the soil by adding tap water.

### Observation and data collection

Evaluation of salinity tolerance was carried out at the reproductive stage based on the following agro-morphological characteristics: days to heading date (HD), grain yield (GY), shoot dry weight (SW), plant height (PH), panicle number (PN), spikelet fertility (SF), grain number per panicle (GP), panicle length (PL), panicle fertility (PF), visual scoring of salt injury (VSC), and visual tolerance score (VTSC) which is the reverse value of VSC. VSC was performed by the observation of general growth with score 1 (growth and tillering nearly normal) to score 9 (almost all plants dead or dying), in relation to standard resistant and susceptible checks at the stem elongation to booting stages based on the Standard Evaluation System of rice established by IRRI (2013). HD was recorded at 90% of emerged panicles in days from seed germination. In the field, GY was calculated from 16 harvested plants and adjusted to 14% grain moisture. Other traits (PH, PN, SF, SW, GP, and PL) were measured from 5 harvested plants. SF was calculated by the percentage of filled grains within a panicle. In the pot experiment these parameters were recorded from the two plants per pot. Panicle fertility (PF) was recorded additionally in the pot experiment where PF was the percent of panicles producing seeds.

### Statistical analysis

Means, standard deviations and analysis of variance (ANOVA) were considered for all measured parameters to determine significant effects (P<0.05) of salinity treatments, genotypes and their interaction. To account for block effects in the augmented RCDB and explore differences between genotypes within salinity treatments, the data was further analyzed using the augmentedRCDB package in R [33]. Then, Pearson’s correlations, Stepwise Linear Regression, PCA and Cluster analysis were performed by using R software. PCA and Cluster analysis were performed under FactoMineR (Factor analysis and data mining with R) package [34].

## Results

During the vegetative stage until maximum tillering, salinity was kept low by frequent flushing/irrigation and EC values remained around 0.34 dSm^-1^ in both low and severe salinity fields (Table 1). The frequency of irrigation was reduced during the reproductive stage (from the elongation stage) and that increased EC to 2.22 - 3.50 dSm^-1^ (mean of 2.57 dSm^-1^) in the low salinity field and to 3.61 - 7.20 dSm^-1^ (mean of 4.73 dSm^-1^) in the severely saline field.

**Table 1.**
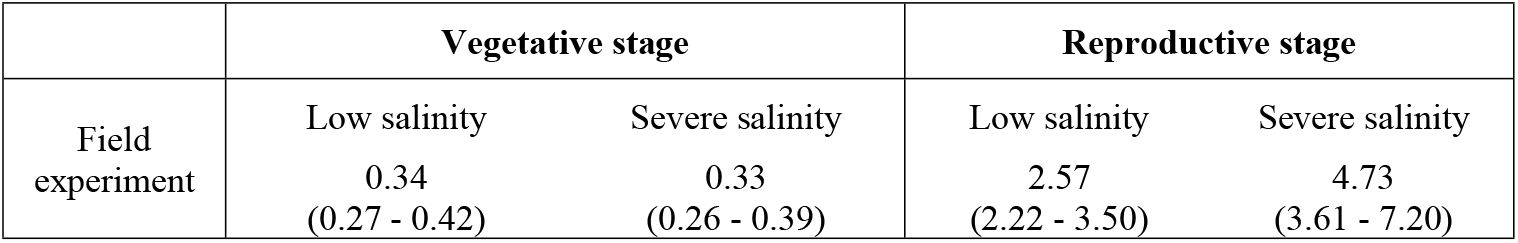
Electric Conductivity (EC in dSm^-1^) in the field experiment during vegetative and reproductive stages. Values shown are averages of bi-weekly measurements (ranges).

### Effect of salinity in the field experiment

A VSC of 2.6 under low salinity (Table 2) indicated that plant growth was nearly normal except for minor occurrences of whitish leaf tips or rolled leaves. In contrast, severe salinity led to reduced plant growth with most leaves being partly whitish and rolled, only a few elongating and this was reflected by an average VSC of 4.7 (Table 2). These visible symptoms of severe salinity were reflected in a reduction of GY to 39.8 g m^-2^ from 362.9 g m^-2^ in low salinity and a corresponding decrease of SF from 82.5% to 29.6%. Strong negative effects were also detected for PN, PL, GP, SW and PH with respective means of 282.4 panicles m^-2^, 19.7 cm, 78.9 grain per panicles, 336.1 g m^-2^, 75.4 cm in low salinity and 76.9 panicles m^-2^, 18.1 cm, 59.8 grain per panicles, 151.2 g m^-2^, 55 cm under severe salinity (Table 2). Increasing salinity furthermore delayed heading by two days (Table 2). Generally, differences in heading did not appear to impact GY and SF.

**Table 2.** Descriptive statistics and p values from an ANOVA based on an augmented RCB design with check varieties IR64 and TsipalaA grown in a field experiment under two levels of salinity.

| Traits | Low salinity |  |  |  | Severe salinity |  |  |  | p value |  |  |  |
| --- | --- | --- | --- | --- | --- | --- | --- | --- | --- | --- | --- | --- |
|  | Mean | Range | SD | CV | Mean | Range | SD | CV | Effect of salinity (%) | Salinity | Genotype |  |
|  |  |  |  |  |  |  |  |  |  |  | Low | Severe |
| VSC | 2.6 | 0.3 - 3.9 | 0.6 | 23.5 | 4.7 | 2.7 - 6.7 | 0.9 | 20.1 |  | *** | *** | *** |
| GY | 362.8 | 189 – 563 | 75.9 | 21 | 39.9 | 2 – 89 | 19.2 | 48.1 | -89.1 | *** | ** | ** |
| PN | 282.4 | 118 – 443 | 81.7 | 28.9 | 77.3 | 10 – 161 | 42.6 | 55.4 | -72.8 | *** | * | ** |
| SF | 82.5 | 44 – 100 | 10.3 | 12.5 | 29.6 | 4 – 54 | 12.9 | 43.4 | -64.1 | *** | * | NS |
| GP | 78.9 | 45 – 114 | 15.3 | 19.4 | 59.8 | 35 – 114 | 13.3 | 22.2 | -24.1 | *** | *** | ** |
| PL | 19.7 | 16 – 23 | 1.1 | 5.5 | 18.1 | 13 – 21 | 1.4 | 8.0 | -8.3 | *** | *** | NS |
| SW | 336.1 | 113 – 901 | 127.4 | 37.9 | 151.2 | 0 – 375 | 68.0 | 45.0 | -55.0 | *** | NS | ** |
| PH | 75.4 | 55 – 91 | 8.3 | 11.0 | 55.0 | 35 – 75 | 6.9 | 12.6 | -27.0 | *** | *** | ** |
| HD | 83.2 | 77 – 98 | 4.8 | 5.7 | 85.4 | 77 – 102 | 6.0 | 7.0 | +2.6 d | *** | *** | *** |

Significant level \*\*\*, \*\*, \*, and NS means P-value < 0.001, 0.01, 0.05, and not significant, respectively.
**VSC:** visual score of salt injury, **GY:** grain yield gm<sup>-2</sup>, **SW:** straw weight gm<sup>-2</sup>, **PH:** plant height cm, **SF:** spikelet fertility, **GP:** grain number per panicle, **PL:** panicle length cm, **PN:** panicle number m<sup>-2</sup> and, **HD:** heading date after sowing, **HI:** harvest index, **SD:** Standard deviation, **CV:** coefficient of variation.

In the low salinity field substantial genotypic variation was observed for all traits with higher variation for SW, PN, VSC and lowest variation for HD and PL. (Table 2). Highest grain yield was seen in MTM16_1 (562.8 g m^-2^) followed by MTM30_2, MTM34_1 and IR55179 with GY of 514.4, 512.1 and 484 g m^-2^ respectively (Fig 1A and S1 Table). The lowest GY was measured for MTM23_2 (189.1 g m^-2^). Two check varieties were repeated 10 times in the augmented design and their GY was significantly different with averages of 330.6 g m^-2^ (SD=56.0) for IR64 and 453.2 g m^-2^ (SD=76.2) for Tsipala-A (Fig 1A). Their respective SF was 86.4 % (SD=4.1) for IR64 and 80.4 % (SD=6.9) for Tsipala-A (S1 Table). Sensitive local variety Sebota 281 displayed the lowest spikelet fertility (43.9 %) and the highest was in MTM34_1 with a SF of 100 % (S1 Table).

**Figure 1:**
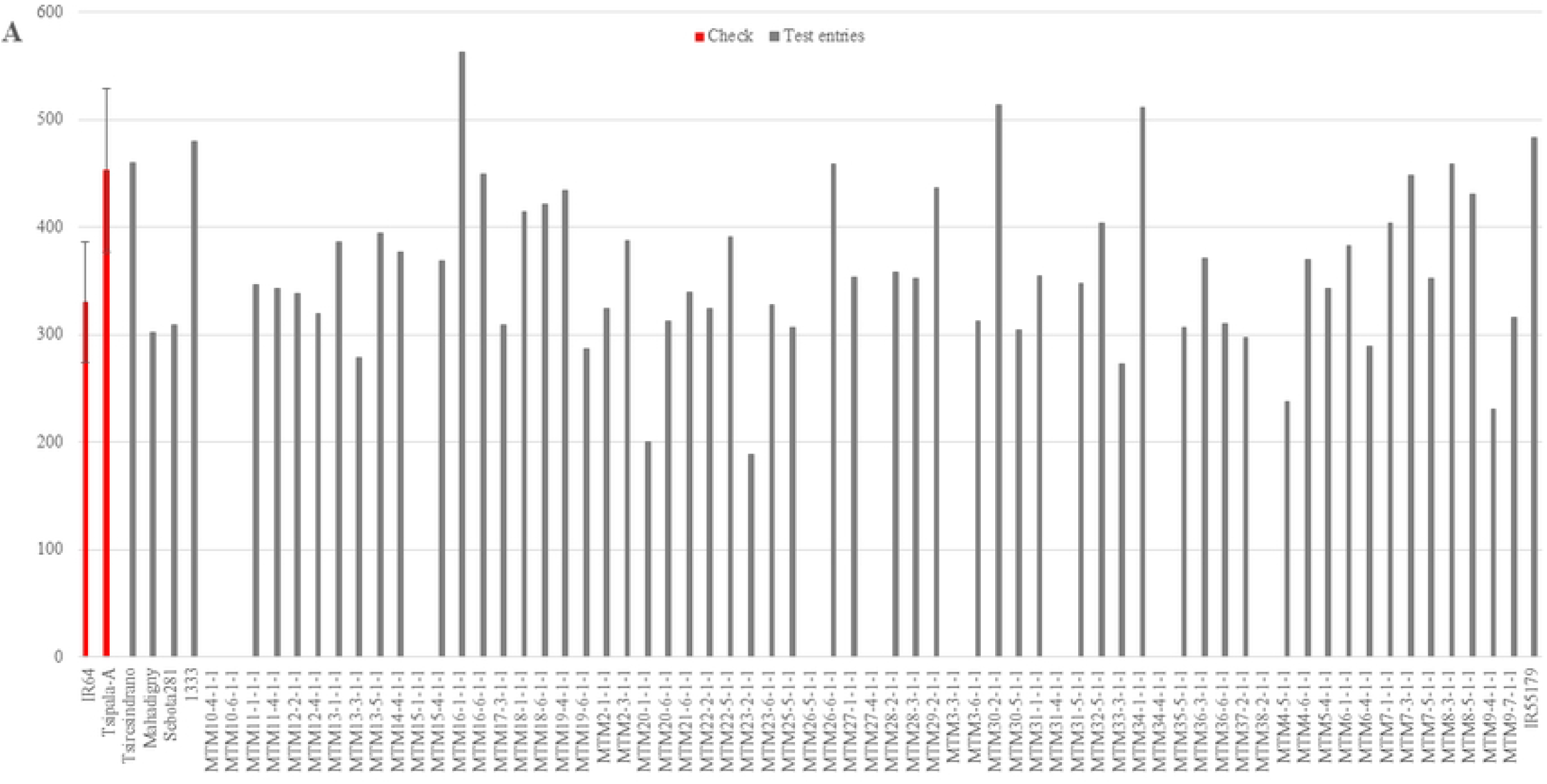
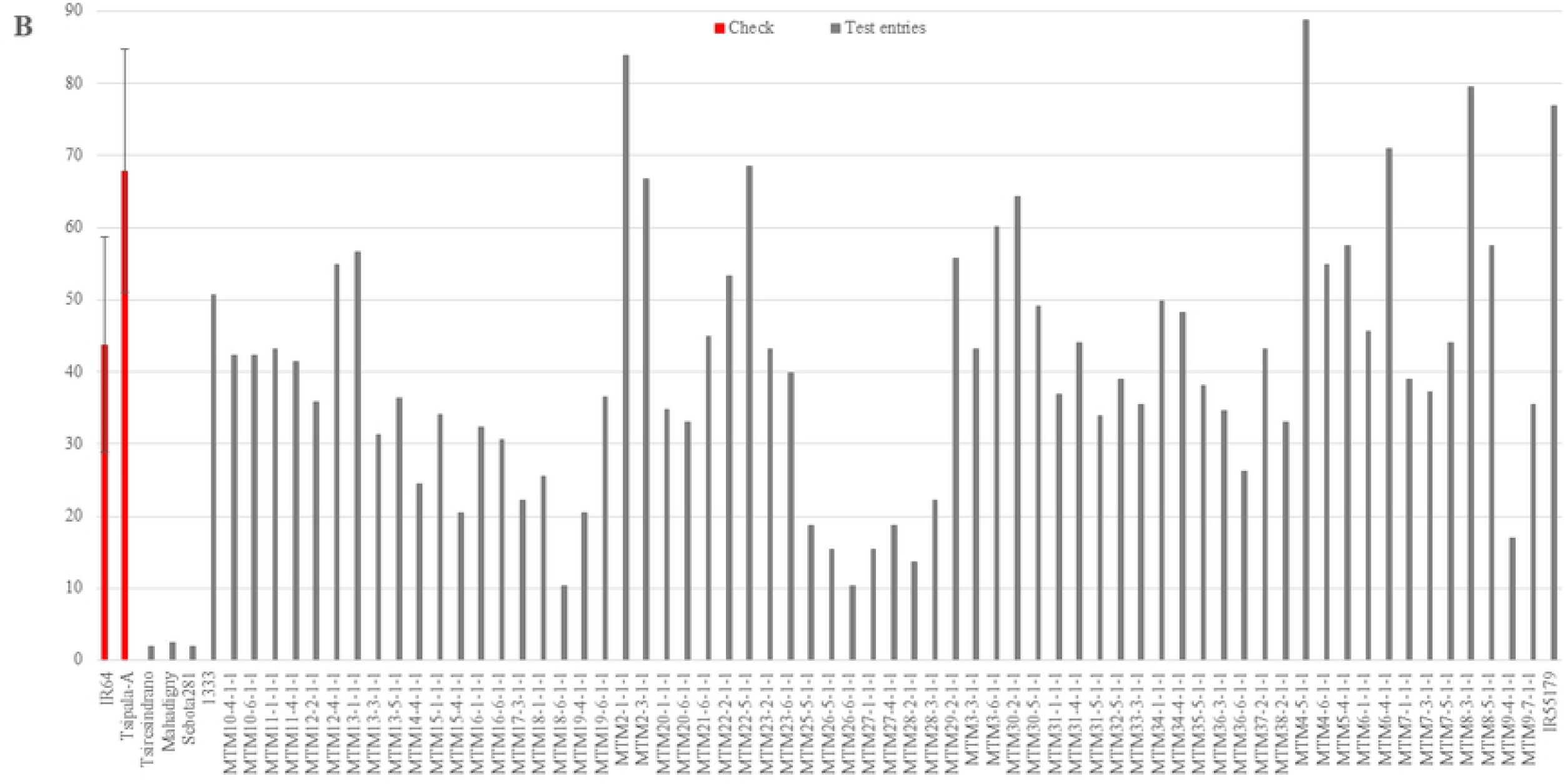
Grain yield (g m^-2^) of checks and test entries from the low salinity (A) and severe salinity (B) fields, based on adjusted values from an augmented RCBD analysis. Bars for checks refer to standard deviations.

Severe salinity drastically affected GY of all genotypes tested and most strongly in the three popular local varieties Sebota 281, Mahadigny and Tsiretsindrano which only yielded around 2 g m^-2^ (Fig 1B) which represents a 99% reduction compared to low salinity. These low yields were accompanied by very low SF of 11 – 15%, compared to an average of 29.6% for all genotypes (Table 2). The two check varieties had higher than average GY under severe salinity and with 67.9 g m^-2^ (SD=16.9) Tsipala-A was significantly better than IR64 with 43.7 g m^-2^ (SD=15) (Fig 1B). Their respective SF was 35.3% (SD=12.1) and 27.5% (SD=10.6) (S1 Table). Check variety IR64 is considered moderately tolerant to salinity and eight of the MTM breeding lines derived from IR64 had a GY above that of IR64 (Fig 1B). One MTM line (MTM4-5) had a GY above the best check (88.8 g m^-2^), then followed by MTM2-1-1-1 MTM8-3, and IR55179 with GY of 83.9, 79.5, and 76.9 g m^-2^ (Fig 1B). IR55179 had the highest SF with 54.2 % and MTM20-1-1-1 had the lowest (3.8 %) (S1 Table).

### Assessment of traits contributing to field salinity tolerance at the reproductive stage

A PCA analysis of all measured traits from severe salinity was conducted, and the first two principal components, which had Eigen values above unity, described 55.6% (PC1) and 14.7% (PC2) of the variation among traits. Traits were separated along PC1 and PC2, with GY, PN, SF, PL, GP, SW and VTSC separating mostly on PC1, while HD and PH contributed to PC2 (Fig 2). GY displayed positive association with PN, SF, GP, PL, SW and VTSC (Fig 2). Using hierarchical cluster analysis and the factor separation provided by the principal component analysis, rice genotypes could be separated into three different salt tolerance groups (Fig 2). Cluster I composed of salt sensitive genotypes including local genotypes Sebota 281, Mahadigny and Tsiresindrano. Salt sensitive genotypes were grouped in the left side of the biplot and presented a low value for GY, PN, SF and GP. Cluster II in the center of the plot included 50% of all lines with intermediate tolerance to salinity. Cluster III on the right of the biplot represented the salt tolerant group with 20% of all genotypes including local salt tolerant variety 1333 and check Tsipala-A. This salt tolerant group had high values for GY, PN, SF, PL and GP (Fig 2).

**Figure 2:**
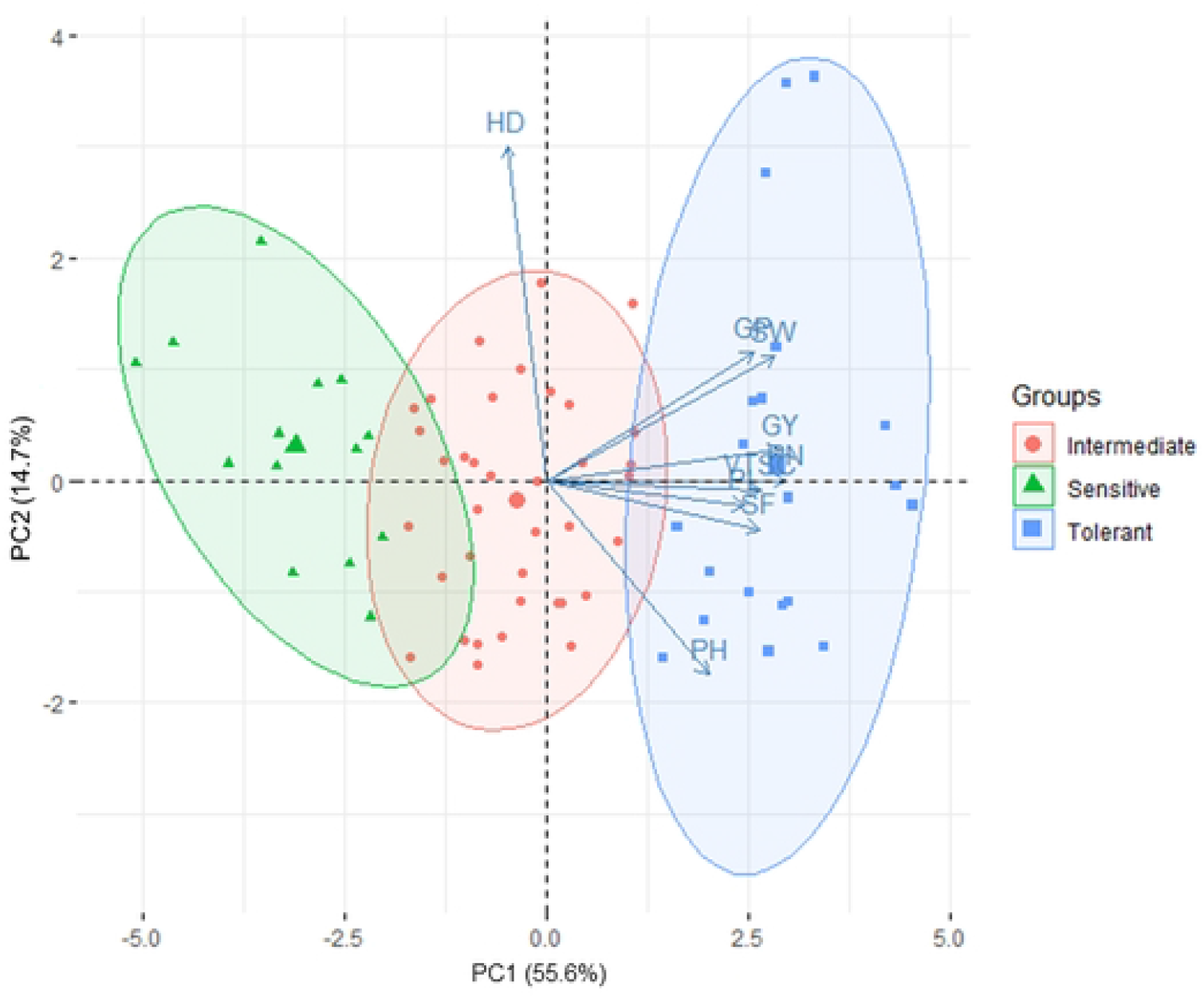
Principal component analysis (PCA) showing the first two principal components (PC1 and PC2) classifying rice genotypes into three different salt tolerance groups based on all agro-morphological parameters measured under severe salinity. Cluster 1: Sensitive, Cluster 2 : Intermediate, Cluster 3 : tolerant, VTSC: visual tolerance score, GY: grain yield per m2, SW: straw weight par m2, PN: Panicle number per m2, SF: Spikelet fertility, GP: Grain number per panicle, PL: Panicle length in cm.

Trait interactions with GY under severe salinity were further analyzed by stepwise linear regression using PN, SF, GP, PL, SW, HD, PH and VTSC as starting variables. The model retained (R^2^ = 0.43) included only PN, SF and GP, with PN being most influential (Table 3).

**Table 3.**
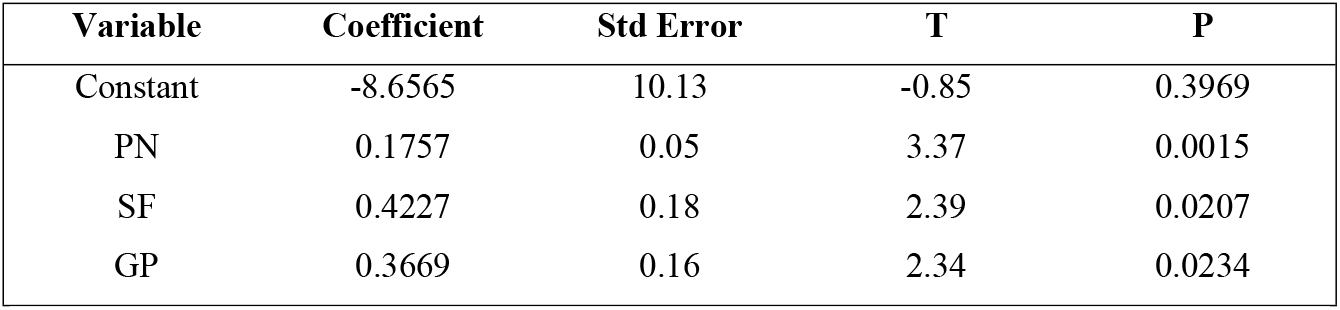
Stepwise linear regression for grain yield as dependent variable and PN, SF, GP as independent variables. PL, SW, HD, PH and VTSC were successively dropped from the model as having a P > 0.05.

### Selection of tolerant genotypes for validation in pot experiment

For further confirmatory studies, eight tolerant genotypes were selected from Cluster III based on a combination of high GY, PN and/or SF (Table 4). Respective values highlighted in bold print are more than one SD above values for parental check IR64. Thus, seven lines were selected for their high PN and four for their superior SF.

**Table 4.** Selected lines for the pot experiment, including the local varieties Tsipala-A (moderately tolerant) and Sebota281 (highly sensitive). The selection of breeding lines was based on having a combination of high GY, PN and SF. Values in bold indicate they exceeded parental check IR64 by more than one SD.

| Genotypes | Low salinity |  |  | Severe salinity |  |  |
| --- | --- | --- | --- | --- | --- | --- |
|  | GY | PN | SF | GY | PN | SF |
| Tsipala-A | 453.2 | 223.0 | 80.4 | 67.9 | 116.0 | 35.3 |
| IR55179 | 484.0 | 312.8 | 89.5 | **76.9** | 73.0 | **54.2** |
| MTM11-1 | 347.1 | 317.8 | 79.7 | 43.1 | **116.1** | 32.3 |
| MTM12_2 | 338.7 | 237.8 | 85.6 | 35.9 | **141.1** | 31.3 |
| MTM13_1 | 386.0 | 257.8 | 84.5 | 56.6 | **106.1** | **38.4** |
| MTM13_3 | 278.7 | 225.3 | 89.4 | 31.3 | **126.1** | 36.1 |
| MTM13_5 | 394.4 | 377.8 | 85.5 | 36.4 | **121.1** | 30 |
| MTM29_2 | 436.7 | 437.8 | 89.6 | 55.8 | **135.5** | **39.9** |
| MTM8_3 | 458.6 | 295.3 | 68.5 | **79.5** | **108.0** | **48.2** |
| Sebota281 | 309.4 | 325.3 | 43.9 | 2.0 | 0.0 | 11.4 |

GY: grain yield per m<sup>2</sup>, SW: Straw weight per m<sup>2</sup>, PN: Panicle number per m<sup>2</sup>, SF: Spikelet fertility,

### Validation - Effect of salinity in pot experiment condition

The pot experiment used non-saline soil in all treatments and an EC of around 0.65 dSm^-1^ was maintained throughout the vegetative phase (Table 5). Addition of salinized water to pots during the reproductive stage increased the EC to 4.20 dSm^-1^ (range 3.00-5.60 dSm^-1^) in the moderate salinity treatment, and to 8.00 dSm^-1^ (range of 6.70-9.10 dSm^-1^) in the severe salinity treatment (Table 5). The moderate salinity negatively affected GY which was reduced by 43.4 %, presumably because SF had decreased to 58.4% from 74.1 % in the control and because GN and PF were significantly reduced by 9.0 and 11.1 %, respectively (Table 6). Increasing salinity further from moderate to severe negatively affected all measured traits with the exception of HD, which remained at 56 days irrespective of salinity treatments (Table 6). GY was reduced by 75.7% relative to the control, and this reduction was largely due to a strong decrease in SF (from 74.1 to 34.8 %) and a reduction in PF (from 100 to 43.1 %). Reductions in GP were the 3^rd^ most influential effect of severe salinity. Genotype x salinity interactions were only significant for the sterility parameters SF and PF (Table 6).

**Table 5.** Electric Conductivity (EC in dSm^-1^) in pot experiments during vegetative and reproductive stages. Values shown are averages of weekly measurements (ranges).

|  | Vegetative stage |  |  | Reproductive stage |  |  |
| --- | --- | --- | --- | --- | --- | --- |
| Pot experiment | Control | Moderate | Severe | Control | Moderate | Severe |
|  | 0.64 | 0.62 | 0.65 | 0.60 | 4.20 | 8.00 |
|  | (0.57- 0.70) | (0.56 - 0.70) | (0.67 - 0.76) | (0.56 - 0.65) | (3.00 - 5.60) | (6.70 - 9.10) |

**Table 6.** Means, ranges, coefficient of variation (%) and p values of analysis of variance for all measured parameters of salinity treatments and among rice genotypes in pot experiment.

| Treatments | HD | GY (g/pot) | SW (g/pot) | PH | SF | GP | PL | PN | PF |
| --- | --- | --- | --- | --- | --- | --- | --- | --- | --- |
| Control | 56.9 a<br>49 – 66<br>9.5 | 51.1 a<br>28 – 76<br>19.0 | 123.0 a<br>74-205<br>25.5 | 102.8 a<br>80 – 127<br>12.0 | 74.1 a<br>56.9 – 85<br>9.8 | 114.6 a<br>61 – 179<br>23.3 | 23.7 a<br>18 – 28<br>10.0 | 22.5 b<br>14 – 33<br>19.6 | 100 a<br>-<br>- |
| Moderate salinity | 56.0 a<br>49 – 66<br>10.8 | 28.9 b<br>15 – 40<br>23.9 | 133.5 a<br>73 – 232<br>26.5 | 100 a<br>70 – 131<br>10.1 | 58.4 b<br>86 – 30.6<br>24.7 | 104.3 b<br>74 – 163<br>19.4 | 23.2 a<br>19 – 26<br>7.7 | 26.5 a<br>11 – 44<br>23.3 | 88.9 b<br>42 – 100<br>14.5 |
| High salinity | 56.5 a<br>49 – 66<br>10.6 | 11.5 c<br>5 – 23<br>35.6 | 102.6 b<br>56 – 169<br>26.7 | 92.5 b<br>69 – 116<br>16.5 | 34.8 c<br>0 – 63.3<br>52.4 | 81.0 c<br>28 – 117<br>25.4 | 21.5 b<br>15 – 23<br>11.5 | 21.9 b<br>13 – 32<br>19.4 | 43.1 c<br>7 – 91<br>5.3 – 91.7 |
| **P value** |  |  |  |  |  |  |  |  |  |
| Salinity treatment | NS | *** | *** | *** | *** | *** | *** | *** | *** |
| Genotype | *** | * | *** | *** | *** | *** | *** | *** | ** |
| Salinity x Genotype | NS | NS | NS | NS | ** | NS | NS | NS | ** |
| P value - genotypes in control | *** | NS | *** | *** | *** | *** | *** | NS | NS |
| P value - genotypes in moderate salinity | *** | NS | *** | *** | *** | ** | ** | NS | NS |
| P value - genotypes in high salinity | *** | ** | *** | *** | *** | *** | * | NS | * |

Significant level \*\*\*, \*\*, \*, and N. S means P-value < 0.001, 0.01, 0.05, and not significant, respectively.
**HD:** heading date after sowing, **GY:** grain yield g/pot, **SW:** straw weight g/pot, **PH:** plant height, **SF:** spikelet fertility, **GP:** grain number per panicle, **PL:** panicle length, **PN:** panicle number per pot in pot experiment and **PS:** panicle sterility.

Selected genotypes included the highly sensitive local Sebota281 and the moderately tolerant local Tsipala-A together with tolerant IR55179 and seven breeding lines derived from the *Pup1* introgression breeding program into an IR64 background (Wissuwa, unpublished). Genotypic differences were more pronounced under severe than moderate salinity (Table 6) and the association between GY and sterility traits SF and PF became very tight (R^2^ > 0.84). IR55179 had the maximum SF and GY with respective means of 54.4% and 15.9 g/pot. Among the breeding lines, MTM 13-5, MTM 13-3 and MTM 13-1 were the most performant with respective SF of 51.6, 44.8, 47.9% and GY of 12.5, 14.6 and 14.1 g/pot. Sebota 281 had the lowest SF (3.9 %) and the lowest GY of only 6 g/pot (Fig 3).

**Figure 3:**
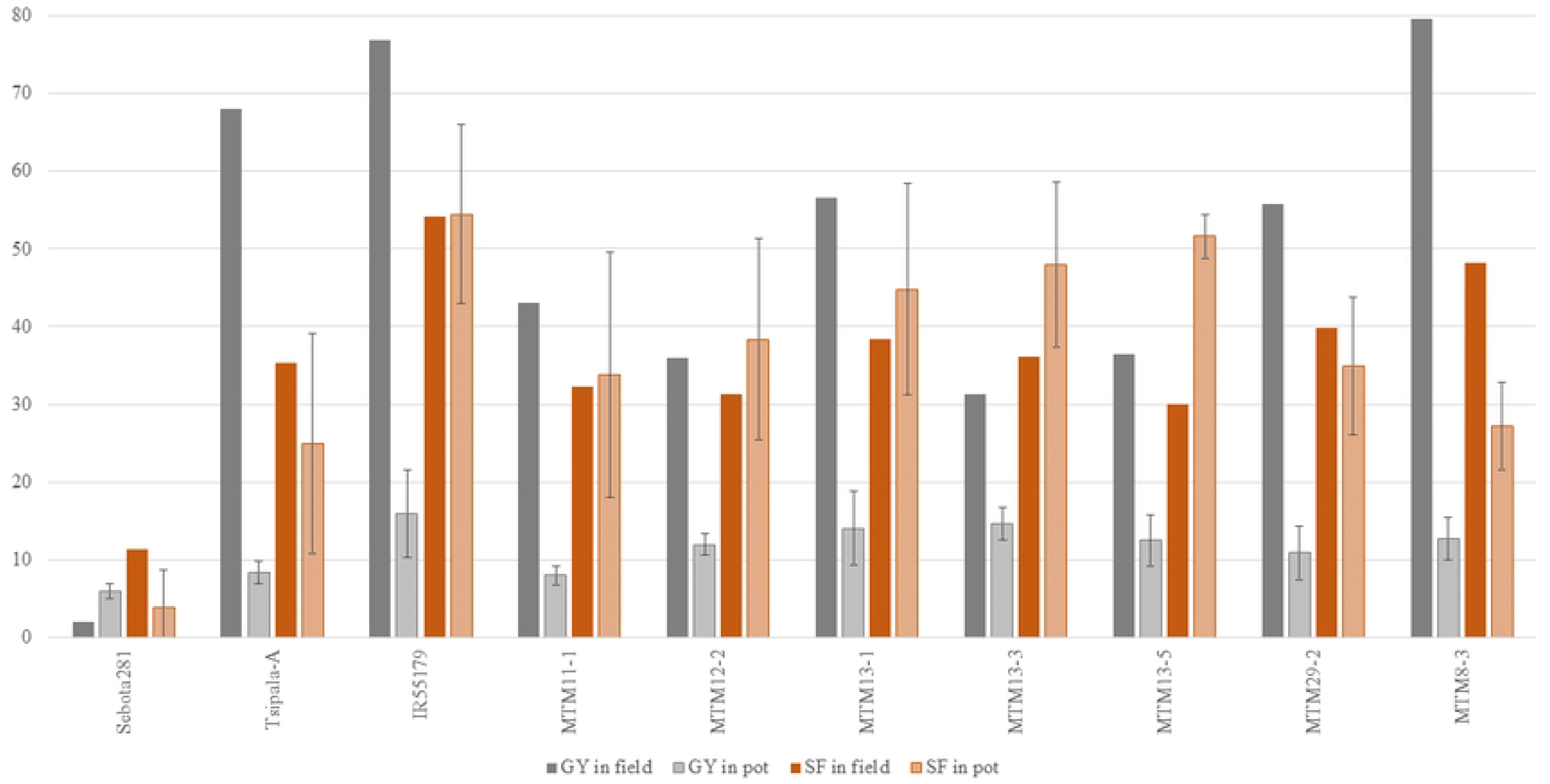
Comparison of selected genotypes in the pot and field experiments for grain yield (GY, in g m^-2^ in field and g pot^-1^ in pot experiments) and percentspikelet fertility (SF). Bars indicate standard deviations.

### Relationship of traits in the pot and field experiment for selected genotypes

Pearson’s correlation of traits from selected genotypes under field and pot experiment showed positive trends (se diagonal in Table 7). Correlations were low for SW and PH, intermediate for GY, PN and GP, and high for SF, PL and HD (Table 7). These finding showed that genotypes response to salinity were similar in the pot and field experiment conditions. Our data confirmed that genotypes with high GY and SF under severe salinity in field condition remained performant in severe salinity under pot experiment (Fig 3). IR55179 was found to be the most tolerant and Sebota 281 most sensitive to severe salinity in both conditions (Fig 3). Likewise, among the breeding lines, MTM 13-5, MTM 13-3 and MTM 13-1 showed high tolerance to salinity in both conditions (Fig 3).

**Table 7.**
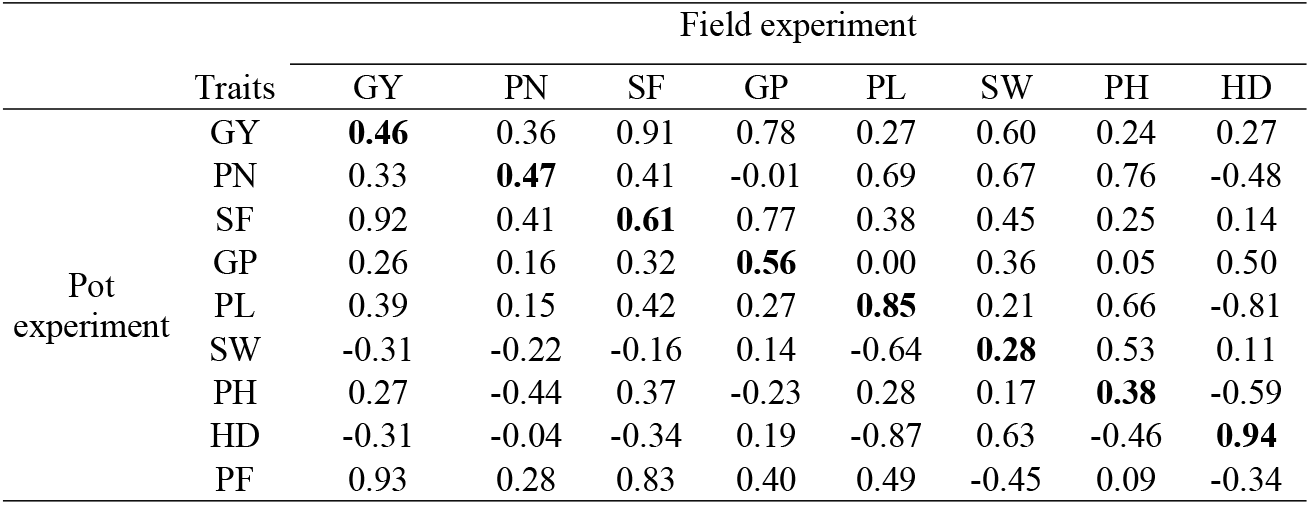
Pearson’s correlation of all traits from field, pot experiment and between field and pot experiment (in diagonal) from severe salinity for the 10 selected genotypes. PF was only measured in the pot experiment.

## Discussion

### Screening for salinity tolerance at the reproductive stage

Rice is highly sensitive to salinity at the seedling and reproductive stages [16]. However, tolerance of salinity at these stages are independent to each other [18]. Plant tolerance at the reproductive stage is very important as it is directly associated with grain yield. Quijano-Guerta and Kirk (2002) [35] reported that the development of salinity tolerant varieties is the most efficient way to address the salinity problem. Thus, it is a prerequisite for rice varieties to possess salt tolerance at the reproductive stage in order to maintain high grain yield in salt-affected regions [36]. In rice, most of the work on salinity tolerance has been conducted at the seedling stage under controlled conditions in the greenhouse using hydroponics or other artificial media, presumably because such methods allow for screening large numbers of lines with relatively high reproducibility [17,37]. As a result of such efforts the major quantitative trait locus (QTL) “*Saltol*” improving salt tolerance at the seedling stage was identified on chromosome 1 from donor Pokkali. Subsequently breeding line FL478 carrying the *Saltol* locus was developed by marker assisted selection (MAS) from a cross of Pokkali and IR29. FL478 has been promoted as an improved donor for the *Saltol* locus for breeding programs to improve salinity tolerance at seedling stage [25,26].

However, limited studies have been conducted on salinity tolerance at the reproductive stage as phenotyping is very complex and difficult. The two major challenges are to manage reproductive stage stress in a way without already injuring plants at the seedling or late vegetative stages, and to impose stress at equivalent growth stages for genotypes that differ in maturity yet are grown together in the same field [30]. In this study, salt stress in the field experiment was avoided at the seedling and vegetative stages by continuously irrigating the field and thereby flushing out any salinity. This technique is commonly used by rice farmers in Marovoay to reduce the effect of salinity in salt-affected fields during the dry season if enough irrigation water is available (Personal communication). Once most genotypes had reached the maximum tillering stage, salinity was induced in the field by reducing the irrigation frequency, thereby allowing the naturally present salt to accumulate in the root zone. In the pot experiment salinity was induced by replacing normal by salinized water. Thus, salinity was limited to the reproductive stage in this study and genotypic differences observed are specific to salt injury during the reproductive stage.

### Detection of genotypes with superior reproductive stage salinity tolerance

Reproductive stage salinity reduced grain yield by 89% on average in the field and by 75.7% in the pot experiments and significant genotype effects were detected in both studies. Pearson’s correlation and stepwise linear regression analysis indicated that maintaining high panicle number and spikelet fertility were the traits contributing most to genotypic differences in yield under salinity at the reproductive stage in the field, while panicle fertility and spikelet fertility were most influential in the pot experiment. Using a principal component analysis combined with hierarchical cluster analysis classified the 72 rice genotypes cultivated under field conditions into tolerant, intermediate and sensitive groups. The tolerant group differed from intermediate and sensitive groups by its higher panicle number, spikelet fertility, panicle length, grain number per panicle, and ultimately, higher grain yield.

The validation experiment conducted in pots with eight selected genotypes from the tolerant group confirmed superior tolerance of IR55179 together with three breeding lines MTM13-1, MTM13-3 and MTM13-5, while Sebota 281 remained the most sensitive. Grain yield was tightly correlated with SF and PF and both traits also showed a high correlation (r = 0.83), possibly suggesting that they are linked to the same physiological tolerance mechanisms. A similarly strong role of percent productive panicles and filled spikelets was reported by Mondal et al. (2019) [38], who also identified QTL associated with these traits. Interestingly, they identified a QTL for grain yield at the same location as a QTL for the number of filled spikelets, but not for the percent filled spikelets, which was mapped to a different chromosomal location.

The effect of salinity on grain yield could have been affected by differences in maturity. Lines used here headed between 76 to 101 DAS in the field and between 49 to 66 DAS in the pot experiment, but differences in heading date did apparently not impact grain yield nor spikelet fertility in our field and pot experiment as indicated by the low correlation of heading with these traits. This is also confirmed by the high tolerance of IR55179 and the three breeding lines MTM13-1, MTM13-3 and MTM13-5, which differed significantly in heading.

The positive correlations between traits in pot and field experiments especially for yield component traits spikelet fertility, grain number per panicle and panicle length, suggest that genotypic responses to salinity were similar under field and pot experimental conditions. Same conclusions were drawn by Kranto et al. (2016) [39] who also report good agreement between field and pot experiments with regard to traits separating tolerant and susceptible rice lines under salinity.

Rice constitutes the main food crop for Malagasy people and is cultivated by most farmers of the country [7]. The north-western coastal region of Madagascar is faced with an increasingly severe soil salinity problem, caused by rising sea levels, increasing temperatures, and changes in rainfall patterns attributed to global climate change [11,40]. Here we could demonstrate that the main rice cultivars grown in the region, Sebota 281, Mahadigny and Tsiresindrano, are highly sensitive to salinity. In order to continuously grow these varieties on salt-affected soil, farmers need to control salinity by flushing salt through frequent irrigation. As water availability tends to decrease towards the end of the dry season when rice is at the reproductive stage, it is feared that salinity will become a growing problem if the locally preferred sensitive varieties continue to be used. Therefore, an urgent need exists to develop tolerant alternatives adapted to the coastal region of Madagascar.

The four tolerant genotypes IR55179, MTM13-1, MTM13-3 and MTM13-5 identified in this study confirm that such tolerant varieties could be identified. While IR55179 is not acceptable locally due to its late maturity and small grains, MTM lines may offer a short-term solution. MTM lines were developed in the IR64 background and while IR64 has never been widely grown in the region, the earlier maturity and taller plant height of MTM lines compared to IR64 would make them locally acceptable. All could potentially serve as donors for improved reproductive stage tolerance, which should ideally be combined with vegetative stage tolerance as conferred by the *saltol* locus.

